# Electron microscopy characterization of minerals formed *in vitro* by human bone cells and vascular smooth muscle cells

**DOI:** 10.1101/661785

**Authors:** Elena Tsolaki, Louis Didierlaurent, Eike Müller, Markus Rottmar, Najma Latif, Adrian H. Chester, Inge K. Herrmann, Sergio Bertazzo

**Affiliations:** Department of Medical Physics & Biomedical Engineering, University College London, London WC1E 6BT, UK; Laboratory for Particles- Biology Interactions, Empa, Swiss Federal Laboratories for Materials Science and Technology, Lerchenfeldstrasse 5, CH-9014, St. Gallen, Switzerland; Laboratory for Biointerfaces, Empa, Swiss Federal Laboratories for Materials Science and Technology, Lerchenfeldstrasse 5, CH-9014, St. Gallen, Switzerland; Magdi Yacoub Institute, Imperial College London, Heart Science Centre, Harefield Hospital, Harefield, Middlesex UB9 6JH, UK

## Abstract

Soft tissue mineralization has been found to be a major component of diseases such as aortic valve stenosis and rheumatic heart disease. Cardiovascular mineralization has been suggested to follow mechanisms similar to those of bone formation with several cell culture models been developed over the years to provide mechanistic insights. These cell models have been characterized by a wide range of biochemical and molecular methods, which identified the presence of osteogenic markers and bone-like cells. However, there is a surprisingly small number of studies where the mineral formed in these cell culture models has been characterized by physico-chemical methods, and even fewer studies have compared this mineral to the one produced by bone cells in cultures. Here we investigated the morphology and composition of the minerals formed in cell cultures of vascular smooth muscle cells and bone cells. Electron microscopy and traditional cell mineralization assays were applied, revealing that vascular cells are indeed able to form calcified nodules of elemental composition similar to bone, however with different morphology. Comparison of morphologies of the two minerals to that found in cardiovascular tissue shows that some of tissue calcification resembles the calcified fibers produced by bone cells *in vitro*. These results suggest that the characterization of the mineral is of utmost importance and its morphology and chemical properties can contribute an important piece of information in the comprehensive analysis of soft tissue mineralization mechanisms, both in *in vitro* cell culture as well as in clinical samples.

## Introduction

Healthy hard tissues, including bone and teeth, are naturally created in the body by mineralizing cells such as osteoblasts and odontoblasts. Mineralization of soft tissues has also been extensively described in the literature and generally, if not always, it has been attributed to pathological processes in humans. Diseases associated with calcification of the vascular system are major contributors to patient morbidity and mortality worldwide [1]. Vascular calcification is a wide spread phenomenon, with a third of all Americans over the age of 45 developing cardiovascular calcification complications [2, 3]. Mineral formation is a hallmark of major diseases including atherosclerosis [4] and coronary heart disease [5], for both of which there is currently no definitive treatment [5, 6]. Calcific cardiovascular diseases are estimated to lead to up to 23.3 million of deaths per year worldwide in 2030 [7].

Despite its clinical importance and major scientific efforts to understand it, the mechanisms that lead to formation of cardiovascular calcification are yet to be understood. Several studies suggest that cardiovascular calcification is an active process [8], even though for many years it was thought to be the direct result of degenerative processes [9]. Works in the literature that associate cardiovascular calcification formation with bone [10, 11] are mainly based on the identification of proteins such as osteonectin and Runx2 in calcific cardiac lesions [10, 12-16]. On the other hand, more recent research on cardiovascular calcification, showed that these calcifications are formed by three distinct forms of mineral; calcified particles, calcified fibers and a compact calcification [1], all of which have a structure significantly different from native bone [1, 17-19].

Cell cultures have been an integral part of cardiovascular research, since they provide a high degree of control of cell environment and allow the induction of calcification [20]. For example, it has been previously shown that the presence of calcium phosphate in cell cultures of mesenchymal stem cells promotes bone formation [21] and that calcification can be induced in vascular smooth muscle cells (VSMC) in the presence of inorganic phosphate [20, 22-24]. Several works suggest that the capability of VSMC to produce calcification *in vitro* [25] is driven possibly by a transdifferention of VSMC to bone-like cells [11, 26, 27]. On the other hand, recent studies demonstrated that the formation of calcification in the vascular tissue or from VSMC can be related to the release of extracellular vesicles [28-30], in a process different to the one observed in bone cells. In light of these seemingly conflicting results, it becomes clear that cardiovascular calcification is a much more complex phenomenon, possibly consisting of a mixture of osteogenic and non-osteogenic processes, and that our current assays and endpoints may not necessarily be sufficient to characterize calcifying cell cultures, as we have yet to understand their underlying mechanisms.

Here, we used physico-chemical methods to characterize the morphology and chemical composition of the minerals present in calcification produced by human bone cell (HBC) and human vascular smooth muscle cell (HVSMC) cultures. We then compared the morphology and composition of the minerals generated *in vitro* to minerals found in human bone and in calcified vascular tissue. Our results show clear structural differences between HBC and HVSMC mineralization, and similarities between the mineralized fibers formed in HBC cultures to mineralised fibers found sometimes in late stage cardiovascular disease.

## Materials and Methods

### Cell culture experiments

Human vascular smooth muscle cells (HVSMC) were acquired from ATCC®. For the different experiments (detailed below), the cells used were kept at -80°C and were used at passage four to eight. Cells were cultivated in vascular cell basal medium (ATCC®) complemented with vascular smooth muscle growth kit (ATCC®).

Bone marrow samples were harvested from patients undergoing surgical hip replacement (ethical approval was obtained from the local ethics committee; EKSG 08/14). The detailed isolation procedure of the human bone cells (HBC) is described elsewhere [31]. Human bone cells (HBC) were cultivated in a proliferation medium consisting in α-MEM (Gibco®), 10% Fetal Calf Serum (FCS), 1% Penicillin-Streptomycin-Neomycin (PSN) and 1 ng/ml basic fibroblast growth factor (FGF-2). Human dermal fibroblast cells (HFC, primary cells, Sigma Aldrich, Buchs, Switzerland) were cultivated in DMEM supplemented with 15% FCS and 1% PSN/L-glutamine. During the cell culture experiments, Lactate Dehydrogenase (LDH) release was measured regularly (twice per week) using a CytoTox 96® Non-Radioactive Cytotoxicity Assay kit (Promega).

For experiments, cells were seeded on 24, 48 and 96 well-plates and grown to confluence. Cells were treated with different media for 28 days. Medium was exchanged every third day.

The following four media compositions were used: (i) α-MEM (*α-MEM*): containing only α-MEM (ii) Proliferation medium (*Prolif*): α-MEM, 10% FCS, 1% PSN and 1 ng/ml basic FGF-2. (iii) Differentiation medium (*Diff*): α-MEM, 10% FCS, 1% PSN, 10 nM 1.25 dihydroxy-vitamine D3 (VitD3), 50 μM ascorbic acid phosphate, 2 mM β-glycerophosphate and 10 nM dexamethasone (Dex) (iv) Differentiation medium + Calcium (*DiffCa*): α-MEM, 15% FCS, 1% PSN, 6 mM calcium chloride, 10 mM sodium pyruvate, 10μM insulin, 50μg/ml ascorbic acid, 10 mM β-glycerophosphate and 1μM Dex. To exclude any influence of compromised cell viability, lactate dehydrogenase content was monitored over the full duration of the experiments and cell viability was measured by MTS assay at day 28.

### Scanning Electron microscopy (SEM)

For SEM imaging, samples were fixed in 4% (w/v) formaldehyde, dehydrated through a graded ethanol (Sigma, ACS reagent 99.5%) series (20, 30, 40, 50 70, 80, 90, 100 and 100% (v/v)) and air dried. All samples were carbon coated and silver painted. A Hitatchi S-3499N, a Carl Zeiss VP and a LEO Gemini 1525 FEGSEM were used for SEM analysis, which involved imaging of the samples using secondary electron (SE) and backscattering electron (BSE) modes, for density-dependent colour SEM (DDC-SEM) to be acquired [1]. Energy Dispersive X-ray Spectroscopy (EDS) analysis was also carried out using Oxford Instruments EDS detectors, integrated into the microscopes. For the analysis, accelerating voltages of 5 kV and 10 kV were used.

### Alizarin Red staining

Following paraformaldehyde fixation, cells were stained with Alizarin Red S to reveal macroscopic calcifications. Alizarin Red S (2 g) was dissolved in 100 mL of distilled water and pH was adjusted to 4.2. Cells were stained with freshly prepared Alizarin Red solution for 2-3minutes.

### Calcium Assay

After the samples were washed three times with PBS, cells were lysed in 1 M HCl for 3 h at 37 °C under constant agitation. Calcium concentrations in the solutions were measured using a colorimetric QuantiChrom™ Calcium Assay Kit (BioAssay Systems) following the protocol provided by the manufacturer. In Brief, 5 μl of each lysate was transferred to a 96-well plate and 195 μl of working reagent (Quanti ChromTM Calcium Assay DICA-500, Gentaur) were added. After 3 min of incubation, the absorbance of the obtained solution was measured at 595 nm (Biotek Instruments Elx 800, Witec AG). All samples were analysed in triplicates and calcium concentrations were calculated by means of a calibration curve using a calcium standard.

### Human sample procurement and preparation

All samples were collected under the guidelines of ethical approval, which included informed consent that allowed us to anonymously analyse the tissues. Human femoral head bone samples were obtained by A. Hart from Charing Cross Hospital (London) from patients undergoing elective total hip arthroplasty procedures. Aortic valves from thirty-two patients were provided by the Oxford Heart Valve Bank at John Radcliffe Hospital (Oxford) and the Royal Brompton Hospital (London). These samples were obtained either after having been rejected for use as homografts or from patients undergoing aortic valve replacement surgery.

All samples were fixed using 4% (w/v) formaldehyde (Sigma, BioReagent, ≥36.0%) solution in phosphate buffered saline (PBS; Sigma) at room temperature immediately after collection, for at least one day.

## Results and Discussion

We investigated the mineralization of HBC and HVSMC in different cell culture media. Pure α-MEM (α-MEM) and proliferation medium (*Prolif*) was used as base line (negative control). Mineralizing media used by representative literature studies were included in order to study *in vitro* mineralization with media containing 2 mM β-glycerophosphate (*Diff*) [31], and media supplemented with 6 mM CaCl_2_ and an additional 8 mM β-glycerophosphate (*DiffCa*) [32].

Total calcium concentration was measured in the cell cultures following 28 days of culturing with media changes every third day (Fig. 1). HBC accumulate significant amounts of calcium irrespective of the mineralizing medium (*Diff* and *DiffCa*), while HVSMC only show an increase in calcium concentration when exposed to calcium-supplemented mineralization medium (*DiffCa*). Interestingly, human fibroblast cells (HFC), which were included as non-vascular cells for reference, show similar calcium accumulation to the other cell cultures when exposed to *DiffCa* for 28 days.

**Fig 1.**
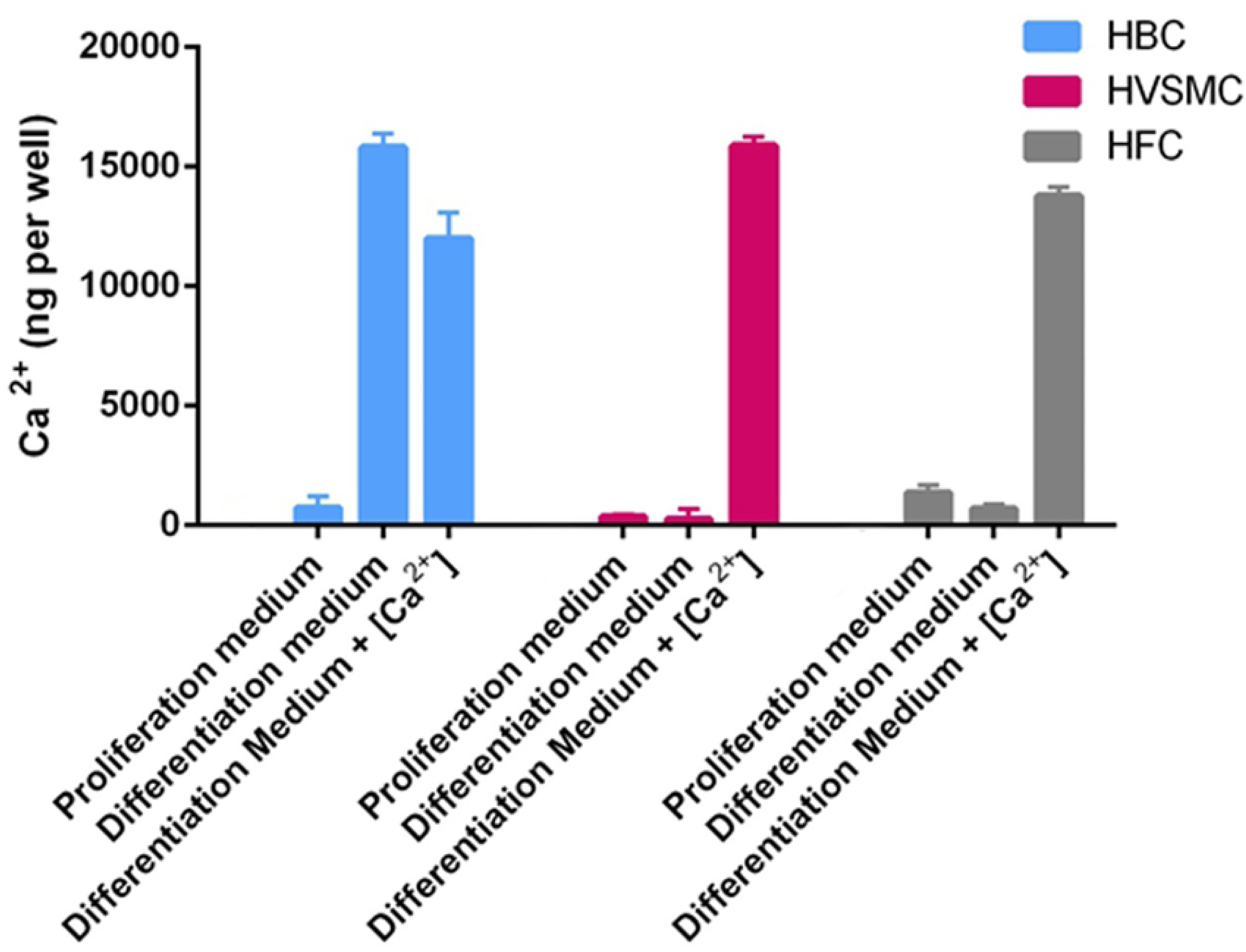
Total calcium content per well at day 28. Cultures of human bone cells (HBC), human vascular smooth muscle cells (HVSMC) and human dermal fibroblasts (HFC), using different media.

Alizarin Red staining was performed, in order to visualize calcium accumulation in the cell cultures (Fig. 2). The results confirm mineral deposition in HBC cultures when exposed to *Diff* and *DiffCa*, and in HVSMC (and HFC, not shown) cultures, only when exposed to *DiffCa.*

**Fig 2.**
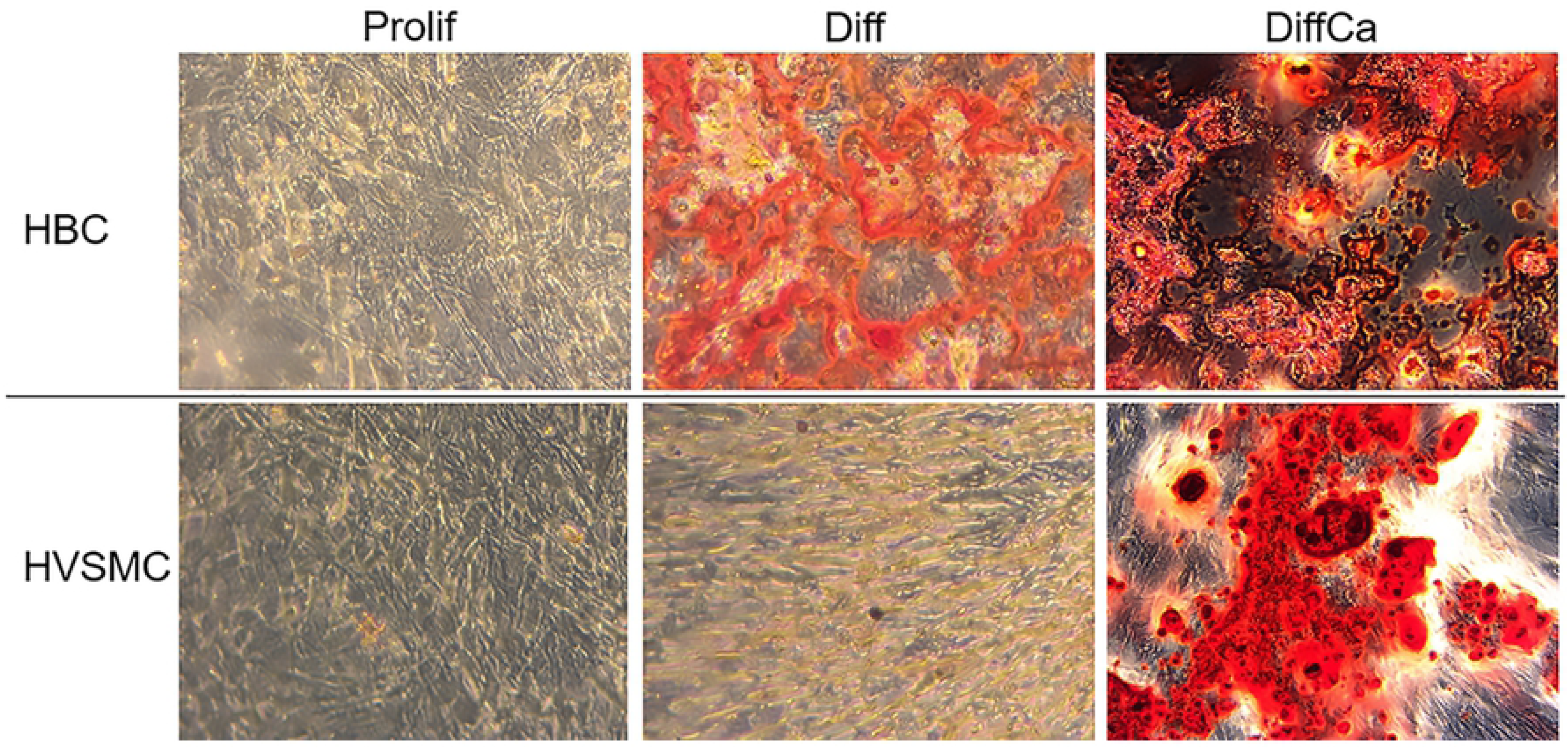
Alizarin Red stained cell cultures. Calcium-rich regions appear in red. Magnification (40x).

Considering that Alizarin Red results may be misleading when applied to cell cultures and can stain positive even in the absence of mineral [33], we also performed high-resolution electron microscopy with elemental analysis. As expected, no evidence of calcium phosphate deposition was found in the HBC culture when exposed to α-MEM and *Prolif* medium *(Fig. 3(a, b)).* In agreement with the calcium assay and the Alizarin Red staining, HBC exposed to *Diff* and *DiffCa* presented significant amounts of mineral deposits. Scanning electron micrographs show fibre-like mineral deposits consisting of calcium phosphate *(Fig. 3(c, d))* in HBC incubated with *Diff* medium. Strikingly, the HBC exposed to *DiffCa* also present large amounts of calcium phosphate mineral deposits *(Fig. 3 (e, f))*, albeit with a completely different morphology to the mineral formed in the *Diff* medium. In the case of *DiffCa*, large chunks of mineral were detected with no evidence of fibrous structures, suggesting a rapid mineral precipitation. While nominal amount and the composition of mineral was very comparable in these two cell culture systems, the morphology of the mineral deposits is strikingly different (see *Fig. 3(c, d) vs. (e, f)*).

**Fig 3.**
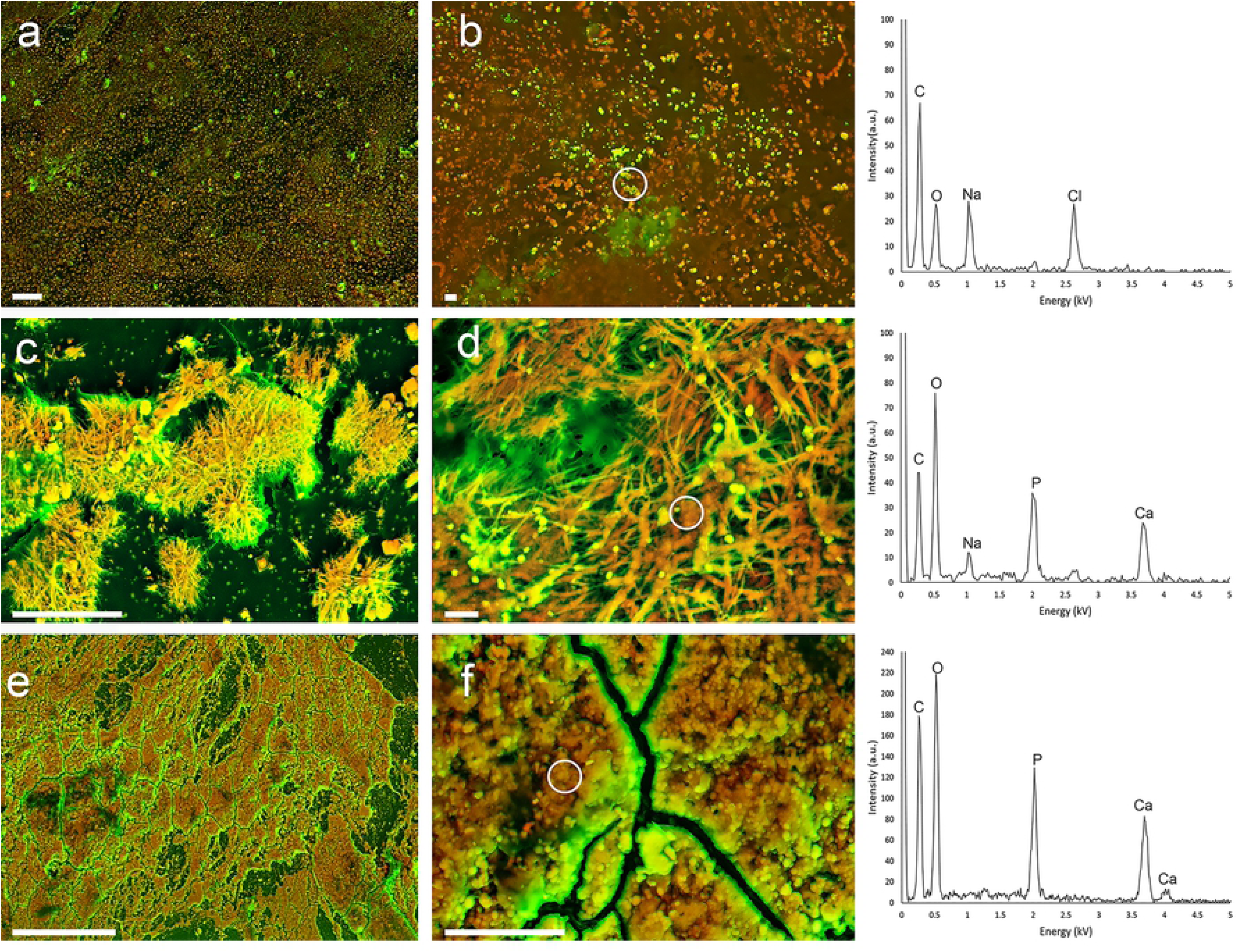
DDC-SEM micrographs of HBC (Human Bone Cells) calcification morphology produced in different media, imaged at low and high magnifications, with correspondent EDS spectra of indicated region. The organic material is represented as green while the inorganic material as orange. (a, b) *Prolif* medium where no deposition of calcium phosphate is observed. (c, d) *Diff* medium where fibrous calcium phosphate structures are observed. (e, f) *DiffCa* medium where large deposition of calcium phosphate is observed. Scale bars = 20 μm, 2 μm.

For HVSMC, again, no evidence of calcium phosphate mineral deposition was found for cell cultures exposed to α-MEM and *Prolif* medium *(Fig. 4 (a, b)).* In contrast to HBC, no evidence of calcium phosphate formation was found for HVSMC exposed to *Diff (Fig. 4 (c, d))*. Calcium phosphate minerals were, however, observed in HVSMC cultures exposed to *DiffCa (Fig. 3 (e, f)).* Similar results were also observed in the HFC culture with mineralization only present in the *DiffCa* medium (results not shown). *DiffCa* medium is indeed saturated for hydroxyapatite [34], and thus it comes as no surprise that rapid mineral precipitation could form in an unstructured manner, similarly to the precipitate formed in the HBC. However the structures observed by the different cell cultures, are significantly different thus a cell involved rather than a simple supersaturation process might be taking place (see *Fig. 3 (e, f) vs. Fig.4 (e,f)*).

**Fig 4.**
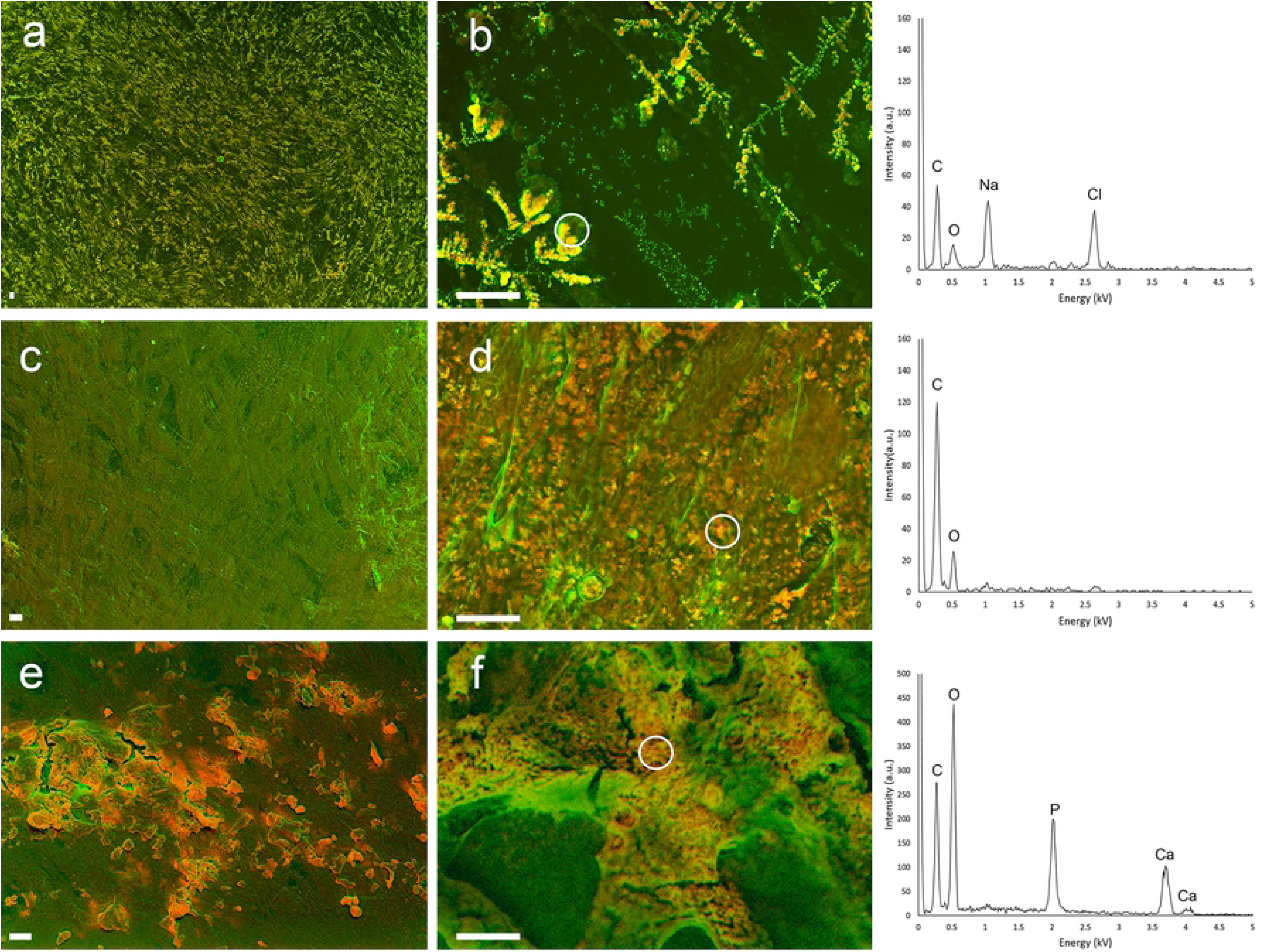
DDC-SEM micrographs of HVSMC (Human Vascular Smooth Muscle Cells) calcification morphology produced in different media, imaged at low and high magnifications, with correspondent EDS spectra of indicated region. The organic material is represented as green while the inorganic material as orange. (a, b) *Prolif* medium where no deposition of calcium phosphate is observed. (c, d) *Diff* medium where no deposition of calcium phosphate is observed. (e, f) *DiffCa* medium where calcium phosphate mineral is observed. Scale bars = 20 μm, 10 μm.

In order to compare the structure and morphology of the *in vitro* generated minerals to minerals found in clinical samples, we analysed a series of calcified human tissues. First, we imaged human bone samples by SEM. The characteristic calcified oriented and regular structure can be easily recognized on the micrographs (*Fig. 5 (a)*). On the other hand, at least three distinctly different characteristic structures, including fibers *(Fig. 5 (b))*, calcified particles *(Fig. 5 (c))* and compact calcification *(Fig. 5 (d))* could be identified in human cardiovascular tissues as previously described in the literature [1].

**Fig 5.**
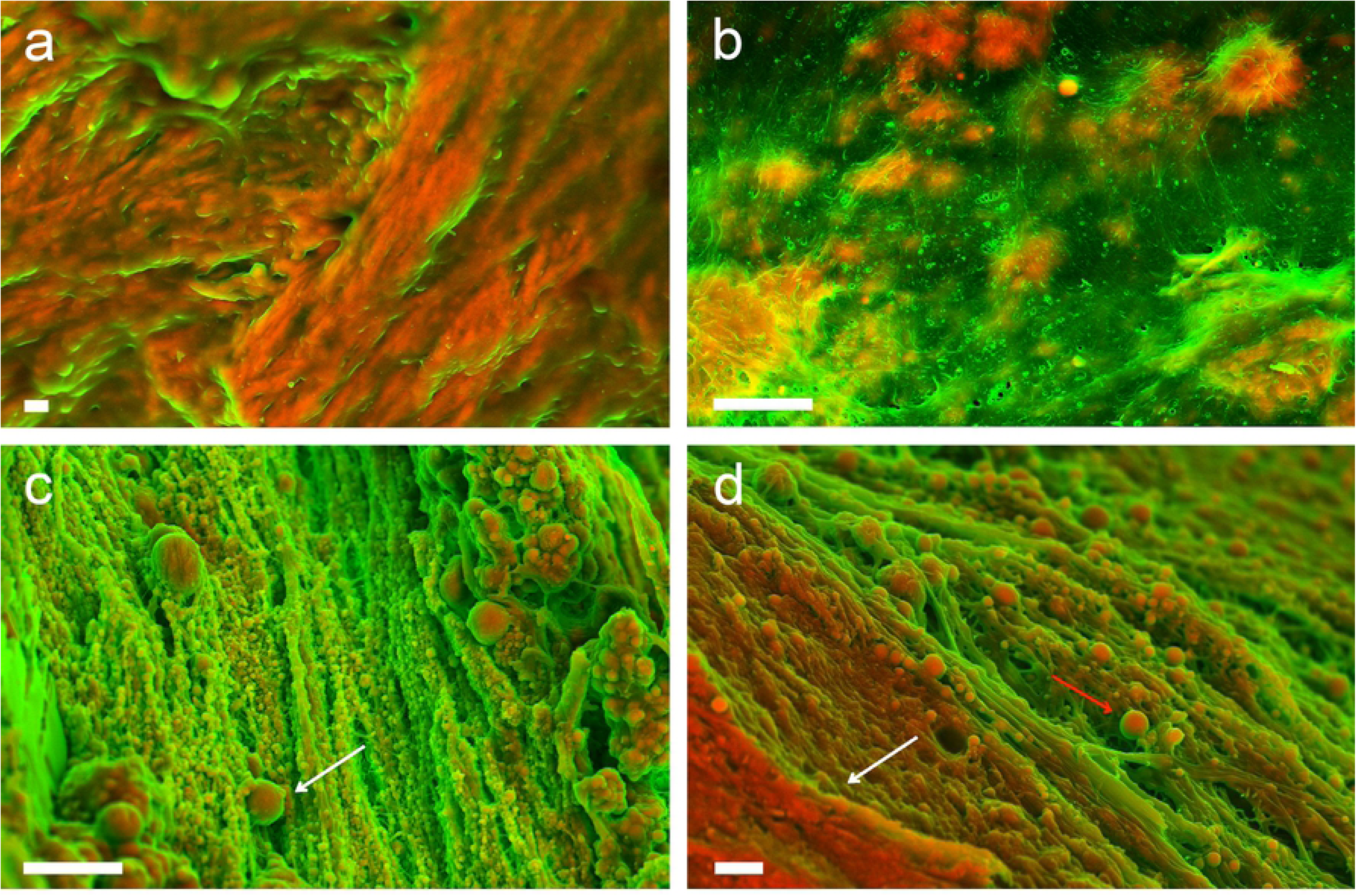
DDC-SEM micrographs of the microstructure of human bone (a) and mineralised structures observed in cardiovascular tissue (b, c and d). (a) High magnification image, showing calcified collagen fibers forming bone. Scale bar = 1 μm. DDC-SEM micrographs of the calcified structures observed in human cardiovascular tissue (b) calcific fibres, (c) calcific particles (white arrow), (d) large compact calcification (white arrow) with calcific particles (red arrow). Scale bars = 1 μm.

There is a notable similarity between minerals formed in the *in vitro* cultures of HBC in *Diff* media *(Fig. 2 (c, d))* and the fibers that can be sometimes found at the later stages of calcification [1] present in the vascular tissue *(Fig. 5b and S1 Fig.).* From the similarities presented between the bone cell cultures calcific structures and the vascular structures, we could speculate that in some cases there is indeed some form of calcification present in the cardiovascular tissue which is similar to the material produced by bone cells, supporting partially the many studies that report osteogenic differentiation of vascular cells. However, no similarities are observed with the natural bone structure, suggesting that the calcified fibers could be formed by a distinct mechanism to bone mineralization. Also, no morphological similarities were found for the mineral produced by HVSMC, neither to the mineral found in bone tissue nor in cardiac tissues, suggesting that our HVSMC model does not adequately depict the *in vivo* mineralization mechanism.

Taken together, these results show that the culturing of human bone cells is indeed able to form minerals similar to some of the structures found *in vivo*, and hence partially support the hypothesis that some sort of osteogenic differentiation in vascular tissue could happen sporadically. On the other hand, none of the cultures were able to produce any structure similar to the calcified particles that are widespread in vascular tissue [1, 17, 30] and are the first calcified structure that can be detected in those tissues [1] (*Fig. 5 (a)*). This study indicates that a major component of vascular calcification still cannot be described by the cell cultures tested here, and that there is likely an unaccounted player contributing to mineral deposition in the cardiovascular system.

## Acknowledgement

I.K.H. and S.B. acknowledge support from the Swiss Heart Foundation and the Swiss National Science Foundation (grant no. 173077).

## Supporting Information

**S1 Figure. BSE electron micrographs of calcification observed in human cardiovascular tissue and in the HBC culture exposed to Diff media.** a Low magnification of fibrous calcification observed in human cardiovascular tissue. b Low magnification of fibrous calcification produced in HBC culture exposed to Diff media. Scale bars = 10 μm. c Higher magnification of calcification observed in human cardiovascular tissue where the calcified fibers along with collagen fibers can be observed. d Higher magnification of calcification produced in HBC culture exposed to Diff media. Scale bars = 1 μm.

